# Diverse *w*Mel variants of *Wolbachia pipientis* differentially rescue fertility and cytological defects of the *bag of marbles* partial loss of function mutation in *Drosophila melanogaster*

**DOI:** 10.1101/2021.01.15.426050

**Authors:** Jaclyn E. Bubnell, Paula Fernandez-Begne, Cynthia K.S. Ulbing, Charles F. Aquadro

## Abstract

In *Drosophila melanogaster,* the maternally inherited endosymbiont *Wolbachia pipientis* interacts with germline stem cell genes during oogenesis. One such gene, *bag of marbles* (*bam*) is the key switch for differentiation and also shows signals of adaptive evolution for protein diversification. These observations have led us to hypothesize that *W. pipientis* could be driving the adaptive evolution of *bam* for control of oogenesis. To test this hypothesis, we must understand the specificity of the genetic interaction between *bam* and *W. pipientis*. Previously, we documented that the *W. pipientis* variant, *w*Mel, rescued the fertility of the *bam^BW^* hypomorphic mutant as a transheterozygote over a *bam* null. However, *bam^BW^* was generated more than 20 years ago in an uncontrolled genetic background and maintained over a balancer chromosome. Consequently, the chromosome carrying *bam^BW^* accumulated mutations that have prevented controlled experiments to further assess the interaction. Here, we used CRISPR/Cas9 to engineer the same single amino acid *bam* hypomorphic mutation (*bam^L255F^*) and a new *bam* null disruption mutation into the *w^1118^* isogenic background. We assess the fertility of wildtype *bam*, *bam^L255F^/bam^null^* hypomorphic, and *bam^L255F^/ bam^L255F^* mutant females, each infected individually with ten *W. pipientis w*Mel variants representing three phylogenetic clades. Overall, we find that all of the *W. pipientis* variants tested here rescue *bam* hypomorphic fertility defects with *w*MelCS-like variants exhibiting the strongest rescue effects. Additionally, these variants did not increase wildtype *bam* female fertility. Therefore, both *bam* and *W. pipientis* interact in genotype-specific ways to modulate female fertility, a critical fitness phenotype.

## Introduction

*Wolbachia pipientis* is a maternally inherited endosymbiotic bacteria that infects over 65% of insect species and manipulates reproduction in a myriad of ways in order to ensure its transmission through the female germline (Werren et al. 2008; Lindsey 2020; Ote and Yamamoto 2020). The phenotypes *W. pipientis* induces in its hosts include cytoplasmic incompatibility (CI), in which embryos of matings between infected males and uninfected females die; male killing, in which male embryos die; feminization of embryos; and manipulation of germline stem cell differentiation in order to increase female fertility (Werren et al. 2008). While *W. pipientis* manipulates its host to increase its own transmission, some *W. pipientis* also provide the host protection against viruses, increase fecundity, modulate thermal preference, and increase longevity (Dedeine et al. 2001; Chrostek et al. 2013; Arnold et al. 2019; Truitt et al. 2019; López-Madrigal and Duarte 2020; Hague et al. 2020).

Understanding the genetic mechanisms that *W. pipientis* uses to manipulate its host has been of immense interest to both basic and applied fields of study. *W. pipientis* is of particular interest as a control for disease vectors such as mosquitoes due to its ability to sweep through a population (due to CI) and then protect the insect from viruses that can also cause human illness such as Dengue, Zika, and Chikungunya (Moreira et al. 2009; Hoffmann et al. 2011; Dutra et al. 2016; Utarini et al. 2021). However, it has been difficult to perform genetic studies of *W. pipientis* function to understand the mechanisms of these host-microbe interactions since *W. pipientis* is an obligate endosymbiont and cannot be cultured. Over the past few years, multiple groups have utilized bioinformatic and in vitro screens to identify candidate *W. pipientis* loci that modulate *Drosophila* phenotypes (Ote et al. 2016; Le Page et al. 2017). These loci have then been expressed in *Drosophila* as transgenes. Through these methods, the *W. pipientis* genes *cifA* and *cifB* (orthologs in *w*Mel and *w*Pip) that cause CI have been identified and functionally validated in *D. melanogaster* (Beckmann et al. 2017; Le Page et al. 2017). These transgenic tools have been used to further define the CI phenotype (Shropshire et al. 2018), investigate the consequences of genetic variation at *cifA* and *cifB* on CI (Shropshire et al. 2021), and identify other host phenotypes affected by *cifA* and *cifB* (Deehan et al. 2021). Another group has identified the *W. pipientis* TomO locus which is one of likely multiple *W. pipientis* loci that interact with *Sex lethal*, a germline stem cell (GCS) gene that controls GSC maintenance and sex determination (Ote et al. 2016).

Previously, we reported that the *W. pipientis* variant *w*Mel genetically interacts with another *D. melanogaster* GSC gene, *bag of marbles* (*bam*). Infection with the *w*Mel variant rescues the fertility defect of females transheterozygous for the classic *bam* hypomorphic allele (*bam^BW^*) and a classic *bam* null allele (*bam^Δ59^*) (Flores et al. 2015). *Bam* is the switch for GSC daughter differentiation in *D. melanogaster* females and the switch for terminal spermatocyte differentiation in males (McKearin and Spradling 1990; Insco et al. 2012; Ting 2013). Although *bam* function is essential for gametogenesis, and *bam* is both necessary and sufficient for GSC daughter differentiation, we observe that it is evolving under positive selection for amino acid divergence in an episodic manner across the *Drosophila* genus (Bauer DuMont et al. 2007; Choi and Aquadro 2014). We see particularly strong bursts of amino acid changes at *bam* in the *D. melanogaster* and *D. simulans* lineages (Bauer DuMont et al. 2007). This episodic pattern of positive selection is consistent with selective pressures that are present in some lineages and not others and that may come and go over time. This is similar to the nature of infections in natural populations, where *W. pipientis* variants are known to infect populations, but also can be lost and replaced fairly often and on both long and short time scales (Bailly-Bechet et al. 2017).

Our observation that *W. pipientis* and *bam* genetically interact led us to hypothesize that *W. pipientis* may drive the adaptive evolution of *bam* (Flores et al. 2015). Since *W. pipientis* is a known manipulator of reproduction, *W. pipientis* may be in conflict with *bam* and other GSC genes for control of oogenesis. While it may seem favorable to host fitness if *W. pipientis* manipulates germline stem cell regulation to increase reproduction, and thus not present an evolutionary conflict, it may not be beneficial for the host if oogenesis is regulated by *W. pipientis*. For example, *W. pipientis* may promote oogenesis when it is not favorable for the host to reproduce due to environmental or physiological factors. To further understand if *W. pipientis* could be in genetic conflict with *bam* and drive its adaptation, we believe it is necessary to better understand the genetic interaction between *bam* and *W. pipientis*.

Fully sequenced variants of *W. pipientis* that infect *D. melanogaster* have been found to cluster phenotypically and phylogenetically into two distinct groups of clades: *w*Mel-like variants (including *w*Mel and *w*Mel2) and *w*MelCS-like variants (including *w*MelCS, *w*MelCS2 and *w*MelPop) that are estimated to have diverged 80,000 fly generations before present (Richardson et al. 2012; Chrostek et al. 2013). The *w*MelCS-like variants predominated in *D. melanogaster* originally but have largely (although not completely) been replaced by *w*Mel-like variants world-wide in the late 20^th^ century (Riegler et al. 2005). The *w*MelCS-like variants provide stronger viral protection, reach higher intracellular bacterial titers in males, and often shorten lifespan compared to the more benign *w*Mel-like variants that now predominate. Additionally, uninfected *D. melanogaster* and those infected with *w*Mel-like variants have a higher temperature preference than *D. melanogaster* infected with *w*MelCS-like variants (including *w*MelPop) (Truitt et al. 2019).

To assess the specificity of our initial observation that *w*Mel rescues the female fertility phenotype of the *bam^BW^*/*bam^Δ59^* hypomorphic mutant, we made use of a genetic tool for functionally assessing *W. pipientis* variation generated by Chrostek et al. (Chrostek et al. 2013), a set of *w^1118^* isogenic lines individually infected with ten diverse *w*Mel *W. pipientis* variants from two of the *w*Mel-like clades and the *w*MelCS-like clade defined by Richardson et al (Richardson et al. 2012). The ten *W. pipentis* variants that we analyze thus include both phenotypic classes of *w*Mel variants: the *w*MelCS-like variants which are higher titer, reduce host lifespan, and confer higher viral resistance and lower thermal preference in contrast to the more recent *w*Mel-like variants.

Since fertility is a phenotype affected by genome-wide variation and both the *bam^BW^* and *bam^Δ59^* alleles were generated in non-isogenic backgrounds, we sought to control the *Drosophila* genetic background to better isolate the effect of *W. pipientis* variation and *bam* genotype on fertility. We used CRISPR/Cas9 to edit the single amino acid change of the original *bam^BW^* hypomorph (Ohlstein et al. 2000) into the *w^1118^* isogenic background, and also to create a new *bam* null allele in the same genetic background. Additionally, we had previously not been able to determine the phenotype of *bam^BW^*/*bam^BW^* females, as the *bam^BW^* allele was isolated over 20 years ago and then maintained over a balancer chromosome which allowed the mutation carrying chromosome to accumulated recessive lethal mutations. Thus, we also used these new lines to analyze *bam* hypomorph and null alleles as homozygotes and transheterozygotes.

We confirmed that the amino acid replacement results in the *bam* hypomorphic phenotype previously described when expressed over a *bam* null allele in uninfected females (Flores et al. 2015). Interestingly, we further show that uninfected females homozygous for the *bam* hypomorphic allele do not exhibit GSC tumors or show reduced fertility compared to wildtype females. We then assessed the effect of infection by the individual *W. pipientis* variants by crossing the cytoplasmic maternal backgrounds infected with each *W. pipientis* variant into our *w^1118^ bam* hypomorph line thereby maintaining the same nuclear background (except for the *bam* locus).

We find that all *w*Mel *W. pipientis* variants tested here do not increase fertility in the wildtype *bam* background, but all *w*Mel *W. pipientis* variants rescue the fertility of *bam* mutant females, with *w*MelCS-like *W. pipientis* showing the highest rescue effects. Therefore, the fertility rescue of the *bam* hypomorph by *W. pipientis* is not due to an overall increase in fertility modulated by *W. pipientis*, but a specific interaction between *bam* and *W. pipientis* genotypes.

## Materials and Methods

### Fly stocks and rearing

Prior to experiments, we raised fly stocks on standard cornmeal molasses food at room temperature. We used yeast glucose food for fertility assays. During experiments, we maintained crosses and stocks in an incubator at 25°C with a 12-hour light-dark cycle. The lines carrying the classic *bam* alleles, *bam^Δ59^* (null) and *bam^BW^* (hypomorph) are described on Flybase (Thurmond et al. 2019) and in Flores et. al (Flores et al. 2015). The *W. pipientis* infected *w^1118^* isogenic lines used in this study were generous gifts from Luis Texiera and described in (Chrostek et al. 2013). We used CantonS males for the *w^1118^*; *bam^L255F^*/*bam*^null^ and *w^1118^*; *bam*^+^/*bam*^+^ fertility assays. We used *w^1118^* isogenic males for the *w^1118^*; *bam^L255F^*/*bam^L255F^* fertility assays. We verified that males were uninfected with *W. pipientis* by endpoint PCR and qPCR using primers described below. To generate the *bam* hypomorphic lines infected with the ten different *W. pipientis* variants, we crossed females from the *w^1118^*; TM2/TM6 stock infected with each variant to *w^1118^*; *bam^L255F^*/TM6 males.

### Generation of *bam* alleles with CRISPR/Cas9

We engineered the new *bam^L255F^* hypomorph and *bam* null alleles described in this study in the *w^1118^* isogenic background using synthetic gRNAs and Cas9 protein as follows.

### gRNAs

We used the flyCRISPR target finder to choose gRNAs with zero predicted off-targets in the *D. melanogaster* genome (Supplement Table S1). We then ordered synthetic gRNAs (sgRNA) from Synthego.

### Cloning

We generated all PCR products for cloning with NEB High Fidelity Q5 master mix, and gel extracted and purified PCR products using the Qiagen MinElute gel extraction kit. We created the donor plasmid for homology directed repair using the NEB HiFi assembly cloning kit and the pHD-attP-DsRed vector from flyCRISPR (Gratz et al. 2014). For plasmid prep, we used Qiagen plasmid plus midi-prep kit and sequenced plasmids with Sanger sequencing (Cornell BRC Genomics Core). We ordered primers for PCR, sequencing, and cloning from IDTDNA (Supplement Table S2).

### Donor sequences

To generate the hypomorphic *bam^L255F^* allele we designed a single stranded oligo donor (ssODN) that contained the single amino acid mutation of the original *bam^BW^* hypomorphic allele (CTT->TTT) as well as a single synonymous change to the closest preferred codon in order to kill the gRNA site upon homology directed repair. We used 80 bp of homology on each side of our targeted change (Supplement Table S1).

To generate the null *bam^null-In2-3xP3-DsRed^* allele, we amplified 1.5 kb homology arms from the *w^1118^* isogenic line and cloned them into the pHD-attP-DsRed vector from flyCRISPR (Supplement Table S3). As we did not need the attP site in our line, we amplified the 3xP3-DsRed cassette from the plasmid and assembled the homology arms, 3xP3-DsRed, and the original vector backbone with the NEB HiFi assembly kit, thereby removing the attP site.

### Injections

All CRISPR/Cas9 injections were sourced to Genetivision and were done in the *w^1118^* isogenic line. The injection mix contained plasmid or ssODN donor, sgRNAs, Cas9 protein (Synthego), and an siRNA for Lig4 (IDT DNA, Supplement Table S1).

For the *bam^null-In2-3xP3-DsRed^* allele, we screened for the eye color cassette in F1’s in house using a Nightsea fluorescent system (DsRed filters). For the *bam^L255F^* allele, the edited nucleotide change disrupts an AflII restriction site, allowing us to screen F1’s by PCR (Promega GoTaq mastermix) followed by restriction digests with AflII (NEB). We prepared genomic DNA using the Qiagen PureGene kit.

We backcrossed females of all CRISPR/Cas9 mutants to *w^1118^* isogenic males for three generations, and then crossed the mutants to the *w^1118^*; TM2/TM6 line to maintain the *bam* mutants over the TM6 balancer. All CRISPR/Cas9 edits in the lines were confirmed by Sanger sequencing (Cornell BRC Genomics Facility).

### PCR assays to detect *W. pipientis*

To test for the presence of *W. pipientis*, we used the Zymo quick DNA miniprep kit to prepare DNA from three replicate samples each with three female flies. For each sample, we used three different primers for endpoint or qPCR (Supplement Table S1). We used the common *wsp* primers (Flores et al. 2015), a primer pair targeting *DprA*, and a highly sensitive primer pair to the ARM repeat (Schneider et al. 2014) for endpoint PCR, modified here for qPCR) (Supplement Table S2).

### Immunostaining

We performed immunostaining as described in Aruna et al. (Aruna et al. 2009) and Flores et al. (Flores et al. 2015). Briefly, we dissected ovaries in ice cold 1X PBS and pipetted the tissue up and down to improve antibody permeability. We fixed ovaries in 4% paraformaldehyde (EMS), washed with PBST (1X PBS, 0.1% Triton-X 100), blocked in PBTA (1X PBS, 0.1% Triton-X 100, 3% BSA) (Alfa Aesar), and then incubated in the appropriate primary antibody overnight. We then washed, blocked, and incubated in the appropriate secondary antibody for two hours then washed and mounted in ProLong Glass NucBlue for imaging (ThermoFisher). We used an anti-Vasa antibody from Santa Cruz biologicals (anti-rabbit, 1:200) and a goat anti-rabbit secondary antibody from Invitrogen (Alexa Fluor Plus 488 1:500). We imaged ovaries on a Zeiss i880 confocal microscope with 405 and 488 nm laser lines at 40X (Plan-Apochromat 1.4 NA, oil) (Cornell BRC Imaging Core). We analyzed and edited images using Fiji (ImageJ).

### Assays for nurse cell positive egg chambers

We used the same assay we previously described to assess the rescue of the original *bam^BW^/bam ^Δ59^* hypomorphic cytological phenotype by *W. pipientis* (Flores et al. 2015). We dissected ovaries from mated 2-4 day old *bam^L255F^*/*bam^null-In2-3xP3-DsRed^* females for all *W. pipientis* variants and the uninfected control in PBS. Ovaries were fixed in 4% paraformaldehyde (EMS), washed with PBST (1X PBS, 0.1% Triton-X 100), and then mounted in ProLong Glass with NucBlue (ThermoFisher). We imaged ovaries on a Zeiss i880 confocal microscope with a 405 nm laser line at 10X (C-Apochromat 0.45 NA, water) and 40X (described above). We analyzed and edited images using Fiji (ImageJ). Developing ovaries consist of cysts containing differentiated nurse cells which are polyploid and feature large, easily identifiable nuclei. In contrast, GSC daughter differentiation is blocked in *bam* loss-of-function ovaries resulting in cysts filled with small GSC-like cells with small nuclei. To quantify the rescue of *bam’s* differentiation function, we counted the number of nurse cell positive egg chambers (cysts) per ovary for uninfected females and females infected with each *W. pipientis* genotype (as these ovaries are small and underdeveloped, they stay intact during the fixation and washing steps).

### Fertility Assays

We used the following *w^1118^* isogenic *bam* genotypes for the fertility assays: *w^1118^*; *bam*^+^/*bam*^+^, *w^1118^*; *bam^L255F^*/*bam^null-In2-3xP3-DsRed^*, and *w^1118^*; *bam^L255F^*/*bam^L255F^*. Additionally, we performed a small control fertility assay using combinations of the alleles described above and the classic *bam^BW^* hypomorph and *bam^Δ59^* alleles.

We performed the fertility assays for *w^1118^*; *bam*^+^/*bam*^+^ and *w^1118^*; *bam^L255F^*/*bam^null-In2-3xP3-DsRed^* in three batches to reduce technical error from too large of an experiment. We included separate uninfected controls in each batch of *W. pipientis* variants and used these to make statistical comparisons of mean progeny per female.

We performed all fertility assays (except for the small *bam^BW^* and *bam^Δ59^* analysis) as follows (and described by Flores et al. 2015):

We collected virgin females and aged them 2-3 days, only using flies that eclosed within 48 hours of each other to reduce developmental variation. We collected virgin males uninfected with *W. pipientis* and aged them for 2-3 days. We distributed males from different bottles across the female genotypes to control for any bottle effects. We individually crossed virgin females to two virgin males. The trio were allowed to mate for 8 days, and then flipped to new vials. After the second 8 days the trio was cleared. The progeny for each trio was counted every other day to get the total adult progeny per female. Progeny per female is reported in increments by day that reflect the days after the first progeny eclosed. For example, we report total progeny for days 1-9 which are the progeny counted on day 1 of eclosion to day 9 of eclosion.

For the control fertility assay containing the *bam^BW^* hypomorph and *bam^Δ59^* alleles, we collected and aged flies as described above, except the progeny per trio was counted only once at the end of the experiment.

### Egg Laying Assay

We collected virgin females for each genotype and *W. pipientis* variant and CantonS virgin males and aged them as described above for the fertility assays. We allowed each trio to mate for 24 hours on grape juice agar supplemented with yeast before flipping them to a new grape juice agar vial. We counted the eggs laid within 24 hours and repeated this for three days.

### Statistics

For the nurse cell assay, egg laying assay, and fertility assays we used estimation statistics to assess the mean difference (effect size) of nurse cells, eggs, and adult progeny between infected and uninfected lines. All estimation statistics were done using the dabest package in Python (v. 0.3.1) with 5000 bootstrap resamples (Ho et al. 2019). Estimation statistics provide a non-parametric alternative to other statistical tests of the difference of the mean (for example, ANOVA), and allow us to understand the magnitude of the effect of *W. pipientis* variation on the *bam* phenotype. We display the data with a swarm plot that shows all of the data points and either a Cumming estimation plot (for more than two sample comparisons) or a Gardner-Altman plot (two sample comparisons) that shows the effect size for each sample compared to the control and 95% bootstrap confidence interval. In text, we report significance as a mean difference (effect size) outside the 95% bootstrap confidence interval.

### Data Availability

Fly lines and plasmids used in this study are available upon request. The authors affirm that all data necessary for confirming the conclusions of the article are present within the article, figures, and tables.

## Results

### *bam^L255F^* is a hypomorphic *bam* allele

We used CRISPR/Cas9 to recreate the same *bam* hypomorphic mutation as the original *bam^BW^* allele (Ohlstein et al. 2000) and a new *bam* null allele in the isogenic *w^1118^* background. Therefore, we have two new *bam* mutant alleles, *bam^L255F^* and *bam^null-In2-3xP3-DsRed^* in the same genetic background so we can compare wildtype bam fertility, the transheterozygous hypomorph/null fertility (as we did previously with *bam^BW^/bam^Δ59^*) as well as assess the phenotype of a the homozygous *bam* hypomorphic genotype which has never been assessed before.

We evaluated if the *w^1118^*; *bam^L255F^* allele recapitulated the *bam^BW^/bam^Δ59^* mutant phenotypes by assessing the phenotype of a transheterozygous *w^1118^*; *bam^L255F^* hypomorph over a *bam* null allele. Since we wanted all of our alleles in the same genetic background and there were no existing *bam* null alleles in the *w^1118^* isogenic background, we used CRISPR/Cas9 to knock in a 3xP3-DsRed cassette into the second intron of *bam*, which resulted in a *bam* null allele in the same *w^1118^* background marked with a trackable eye marker (*w^1118^*; *bam^null-In2-3xP3-DsRed^*, Fig 1B). Females homozygous for this *bam^null-In2-3xP3-DsRed^* allele exhibit the expected tumorous ovary phenotype with no developing nurse cell-positive cysts and are sterile (Fig 2A, fertility data not shown).

**Fig 1.**
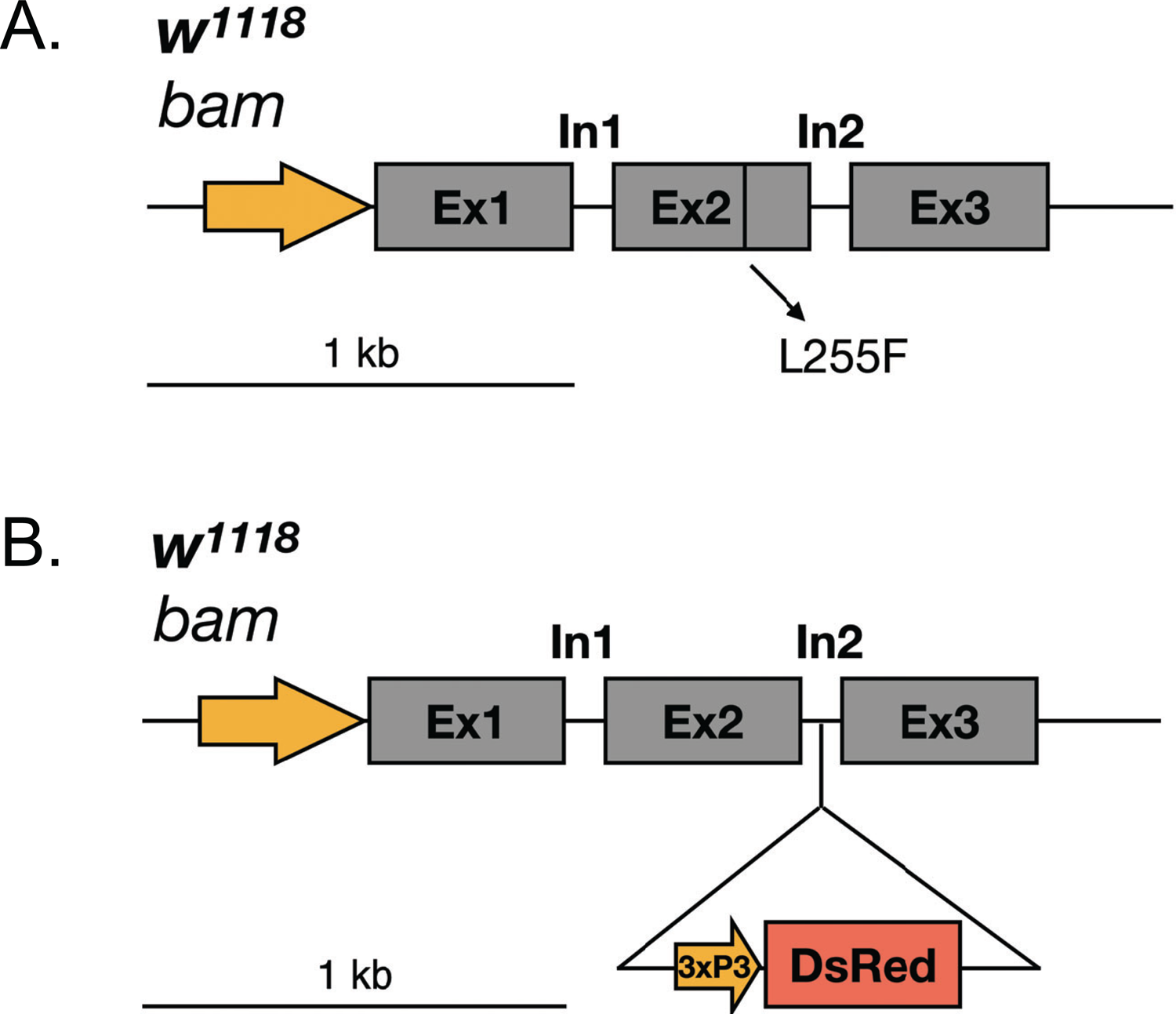
Design for recreating the classic *bam^BW^* hypomorph allele and a new *bam* null allele in the *w^1118^* isogenic background with CRISPR/Cas9 (A) Schematic of the *bam* gene region showing the single gRNA target site in the second exon and the hypomorphic missense mutation. (B) Schematic of the *bam* gene region showing the single gRNA target site in the second intron with the 3xP3-DsRed eye marker to create a *bam* null allele.

**Fig 2.**
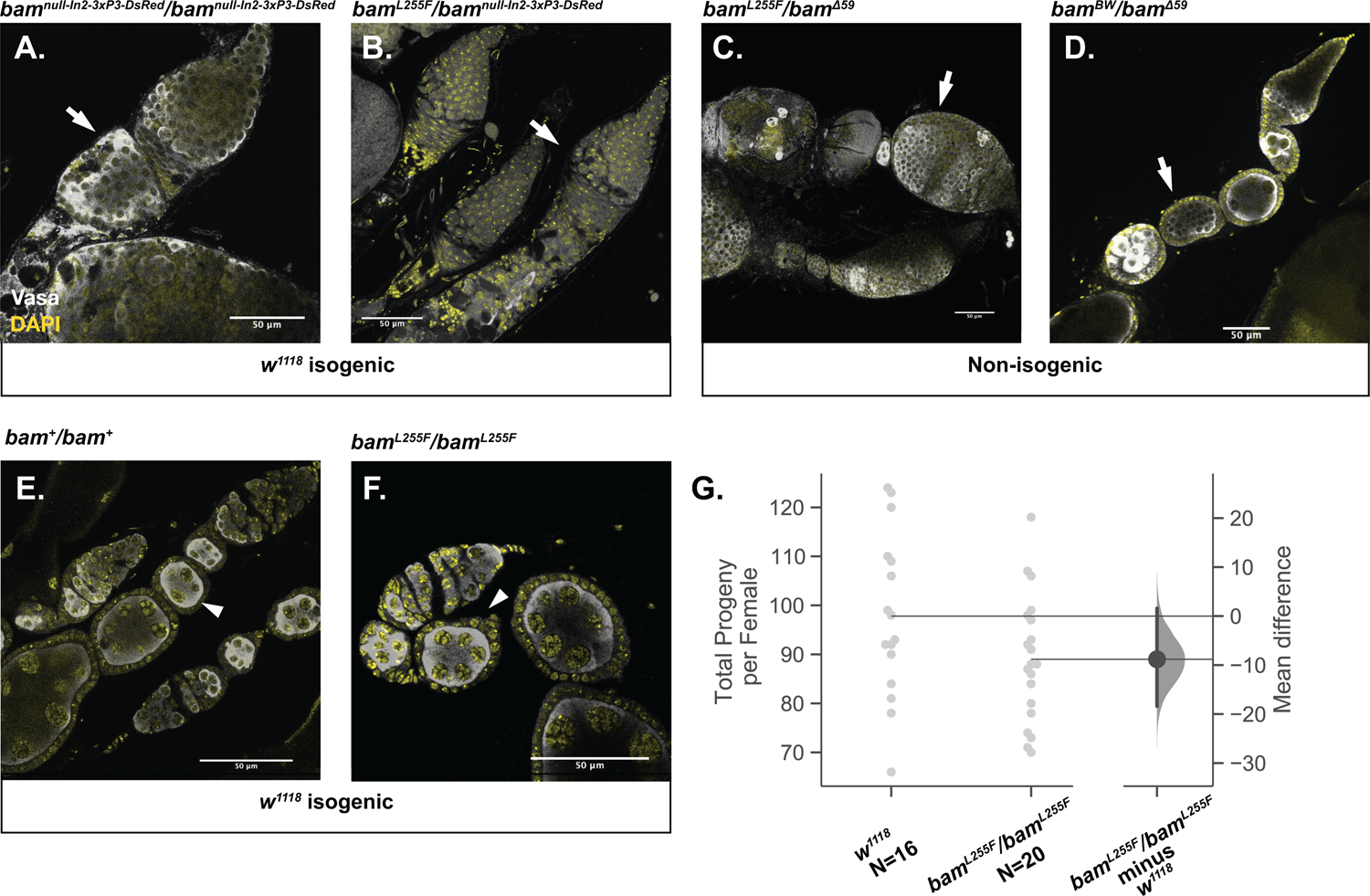
The *bam^BW^* hypomorphic allele and a novel *bam* null allele in the *w^1118^* isogenic background recapitulates the classic *bam* phenotypes. (A) The newly generated *bam^null-In2-3xP3-DsRed^/bam^null-In2-3xP3-DsRed^* genotype results in tumorous ovaries (arrow). (B) The recreated *bam^L255F^* mutation over *bam^null-In2-3xP3-DsRed^* exhibits tumorous ovaries in the *w^1118^* isogenic background (arrow). (C) The recreated *bam^L255F^* mutation over the classic *bam*^Δ59^ null allele exhibits tumorous ovaries (arrow). (D) The classic *bam^BW^/bam^Δ59^* hypomorphic genotype exhibits tumorous ovaries (arrow). (E) Ovaries from the *w^1118^* isogenic line wildtype for *bam* do not exhibit any germline tumors and show developing egg chambers with large nurse cell nuclei (arrowhead). (F) *bam^L255F^*/*bam^L255F^* homozygotes do not exhibit tumorous ovaries, indicating two copies of the partial loss of function *bam^L255F^* mutation is sufficient for GSC daughter differentiation. (G) *w^1118^* females and *bam^L255F^*/ *bam^L255F^* females do not show a significant difference in fertility (mean difference of progeny per female), further indicating that two copies of *bam^L255F^* is sufficient for *bam* function.

We then assessed the phenotype of the *bam^L255F^* allele as a transheterozygous mutant over our new *bam^null-In2-3xP3-DsRed^* in the same *w^1118^* isogenic background. We observed a large fertility defect, but the females were not sterile (Fig 3). The *bam^L255F^*/ *bam^null-In2-3xP3-DsRed^* ovaries exhibited the expected tumorous ovary phenotype, with the presence of some differentiating, nurse cell positive cysts indicating that *bam* is partially functional, and thus that this single amino acid change fully explains the original *bam^BW^* hypomorphic phenotype (Fig 2B). To further verify the nature of the *bam^L255F^* allele, we assessed its phenotype over one of the classic *bam* null alleles that we previously used to study the interaction between *bam* and *W. pipientis, bam^Δ59^*. The *bam^Δ59^* null allele is a nearly full deletion of the *bam* coding sequence. We examined ovaries from *bam^L255F^*/ *bam^Δ59^* females and observed the tumorous ovary phenotype with some developing cysts as has been previously documented for *bam^BW^/bam^Δ59^* and additionally shown here (Fig 2C&D). We measured the fertility of *bam^L255F^*/ *bam^Δ59^* females and observed a large fertility defect, consistent with the fertility of the *bam^L255F^*/ *bam^null-In2-3xP3-DsRed^* genotype (Fig 3). Therefore, the *bam^L255F^*/ *bam^null-In2-3xP3-DsRed^* cytological and fertility phenotypes are consistent with the previously documented *bam* hypomorph phenotypes.

**Fig 3.**
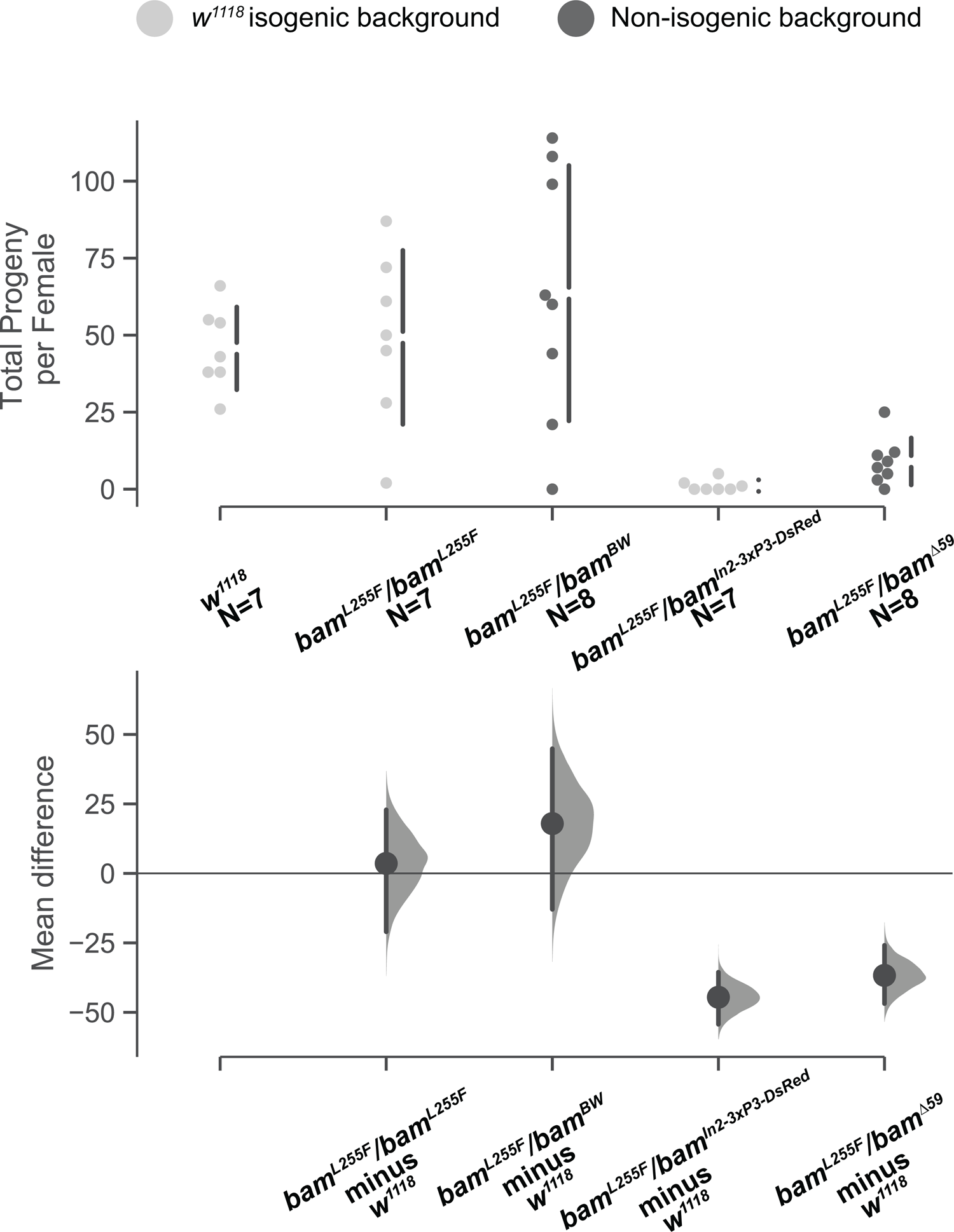
Female fertility of genotypes containing the *w^1118^; bam^L255F^* hypomorphic allele are consistent with those of classic alleles. Swarm and Cumming estimation plots of a control fertility experiment showing that the fertility of *bam^L255F^*/ *bam^L255F^* females is not significantly different from *w^1118^* (mean difference = 3.6), nor are *bam^L255F^*/*bam^BW^* females (mean difference = 17.9). *bam^L255F^*/ *bam^null-In2-3xP3-DsRed^* females have significantly lower fertility compared to wildtype (mean difference = −44.6, bootstrap 95% confidence interval, effect size), as do *bam^L255F^*/*bam^Δ59^* females (mean difference = −36.7, bootstrap 95% confidence interval, effect size).

As the *bam^BW^* allele is homozygous lethal due to accumulated recessive lethal mutations on the third chromosome, we used our *bam^L255F^* allele to ask if the homozygous *bam^L255F^* females also had a hypomorphic phenotype. We asked if *bam^L255F^*/*bam^L255F^* females also exhibited the classic *bag of marbles* cytological phenotype and reduced fertility. In contrast to the tumorous ovary phenotype of *bam^BW^/bam^Δ59^* and *bam^L255F^*/ *bam^null-In2-3xP3-DsRed^*, a result of a defect in *bam*’s differentiation function which causes GSC-like cells to over-proliferate, we found no evidence of tumorous cysts in ovaries of *bam^L255F^* /*bam^L255F^* females (Fig 2F). These ovaries resembled those of a wildtype *bam* background, with all developing egg chambers featuring nurse cells (Fig 2E). Previously we were unable to compare the fertility of the *bam^BW^* hypomorph to wildtype *bam* fertility as fertility is greatly affected by different genetic backgrounds. We compared the fertility of the *w^1118^*; *bam^L255F^* /*bam^L255F^* females to *w^1118^*; *bam*^+^/*bam*^+^ females and found that the mean difference in total progeny per female between the two genotypes was not significantly different (Fig 2G). While we did not measure all aspects of *bam* function of this genotype, the fertility and cytological data here indicate that the *bam^L255F^* /*bam^L255F^* genotype does not cause a severe *bam* mutant phenotype, as we observed previously for the transheterozygous *bam^BW^* hypomorph (*bam^BW^/bam^Δ59^*). Therefore, two copies of the *bam^L255F^* allele are sufficient for germline stem cell daughter differentiation and fertility.

Although we cannot make strong claims about the effect of *bam* genotype on fertility between lines with different genetic backgrounds, we did perform a small control experiment to ask if our new *bam^L255F^* hypomorphic allele over the classic *bam^BW^* hypomorphic allele also show similar fertility to wildtype females as we observed for *bam^L255F^*/*bam^L255F^* females. We observed that *bam^L255F^*/*bam^BW^* females and homozygous *bam^L255F^* females did not show significantly different fertility from *w^1118^ bam^+^/bam^+^* females (Fig 3). Further, we asked if our new *bam^L255F^*/*bam^null-In2-3xP3-DsRed^* genotype showed a similar fertility defect to *bam^L255F^*/*bam^Δ59^* (classic null) and found that both showed severely reduced fertility compared to wildtype (Fig 3). These data further confirm that our new *bam^L255F^* and *bam^null-In2-3xP3-DsRed^* alleles behave similarly to alleles we have used in the past to study the interaction between *bam* and *W. pipientis*.

Since we were able to successfully recapitulate the partial-loss of function *bam* cytological and fertility phenotypes with the *bam^L255F^*/ *bam^null-In2-3xP3-DsRed^* genotype as we have previously described (Flores et al. 2015), but we did not observe the *bam* partial loss of function ovarian cytological phenotype in the *bam^L255F^*/*bam^L255F^* females, we focused primarily on the effect of *W. pipientis* variation on the *bam^L255F^*/ *bam^null-In2-3xP3-DsRed^* female fertility and cytological phenotypes.

For convenience and readability, we will frequently refer to the *w^1118^*; *bam^null-In2-3xP3-DsRed^* allele as *bam^null^* and the *w^1118^*; *bam^L255F^* allele as *bam^L255F^*.

### Infection with the ten different *w*Mel variants we assayed does not increase *w^1118^*; *bam^+^/bam^+^* isofemale fertility

First, to determine if the *w*Mel *bam* hypomorph rescue is a consequence of a general increase in female fertility induced by *W. pipientis*, we assessed the effect of *W. pipientis* variation on the fertility of *w^1118^*; *bam^+^/bam^+^* females of which we have lines infected with ten different *w*Mel variants of *W. pipientis* as described by Chrostek et al. (Chrostek et al. 2013) (Fig 4A).

**Fig 4.**
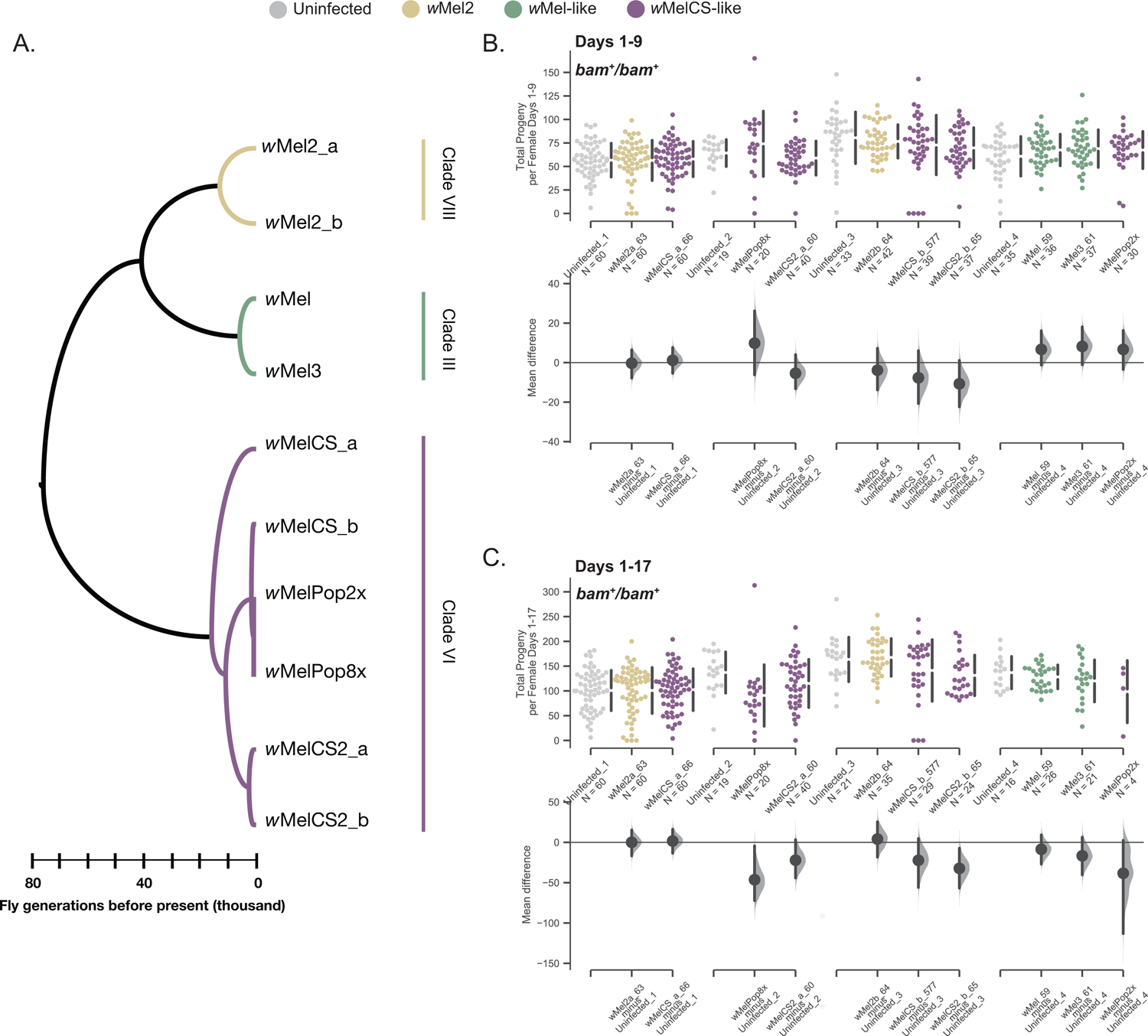
Total progeny per female and mean difference of progeny per female for the *w^1118^*; *bam*^+^/*bam*^+^ genotype infected with ten *W. pipientis w*Mel variants compared to uninfected. (A) Cladogram adapted from Chrostek et al. 2013 showing the relationships and clade assignments for the *W. pipientis* variants tested in this study. (B) Swarm and Cumming estimation plots showing the total progeny per female for each *W. pipientis* infected line counted on days 1-9 post eclosion. As the fertility assays were performed in batches, each batch is compared to its own uninfected control. No *W. pipientis* variant was associated with a significant difference in fertility for wildtype *bam* females over days 1-9 post eclosion (bootstrap 95% confidence interval, effect size). (C) Swarm and Cumming estimation plots showing the total progeny per female for each *W. pipientis* infected line counted on days 1-17 post eclosion. As the fertility assays were performed in batches, each batch is compared to its own uninfected control. *W. pipientis* variants *w*MelPop8X, and *w*MelCS2b were associated with significantly lower fertility over the longer period of days 1-17 post eclosion compared to uninfected controls (bootstrap 95% confidence interval, effect size)

Here, fertility measured over multiple days reflects progeny produced cumulatively by the female over time, although there could also be variation in development time of the progeny. We first assessed the effect of *W. pipientis* variation on wildtype *bam* female fertility (*w^1118^*; *bam*^+^/*bam*^+^). Over 17 days, we counted the progeny produced by each test female infected individually for each of the ten *W. pipientis* variants and assessed the mean difference of adult progeny from uninfected control females. We found that none of the *W. pipientis* variants increased female fertility in the wildtype *bam* background, and some lines showed a significant decrease in total progeny per female across all days measured (*w*MelPop8X, and *w*MelCS2b_60) (Fig 4B&C). Of the lines that showed a negative impact of *W. pipientis* on fertility, this effect was only significant as the female fly aged. On days 1-9 of progeny eclosion, no *W. pipientis* variant had a significantly negative effect on progeny counts, which is notable as these days represent progeny from eggs laid earlier in the test female’s life and thus are more likely to reflect effects of *W. pipientis* variation on fertility in nature (Fig 4B). The variants that showed a significant and large negative effect of *W. pipientis* infection on fertility were both in the *w*MelCS clade (mean difference between −32 and −46 progeny per female for days 1-17 Fig 4C). *w*MelCS variants exhibit higher *W. pipientis* titer in males (Chrostek et al. 2013), which if also true in females may negatively impact female fertility if titer gets too high, as this leads apoptosis of infected cells (Chrostek et al. 2014).

### All *w*Mel variants rescue *bam* hypomorph female fertility but show a range of effect sizes

Our results from the *w^1118^*; *bam^+^*/*bam*^+^ fertility assay indicated that none of the *W. pipientis* variants tested have an overall increase in fertility in females (Fig4). We performed the same fertility assay as described above for the *w^1118^*; *bam^+^*/*bam*^+^ genotype for the *bam^L255F^*/*bam^null^* genotype infected with the same ten *W. pipientis* variants. Females of the *bam^L255F^*/*bam^null^* genotype exhibit tumorous ovaries and are only weakly fertile (Fig 2B, Fig 3). In contrast to the wildtype *bam* lines, in the *bam^L255F^*/*bam^null^* lines we found that all variants increased female fertility both in the first week of progeny eclosion and throughout the entire measured timeframe with the exception of *w*MelPop2X, as infected females died before the end of the experiment, thus we only report the first week of progeny production for this variant (Fig 5A&B). Notably, and some variants had a large effect on female fertility. The variants with the largest effect sizes were in the *w*MelCS-like clade (*w*MelCS2b, *w*MelPop2x, *w*MelPop8x) (Fig 5A&B). This fertility rescue is in contrast to the decrease in fertility by the same *W. pipientis* variants in the wildtype *bam* background, especially for *w*MelCS2b which had the largest negative effect on wildtype *bam* fertility, but the largest positive effect on *bam^L255F^*/*bam^null^* fertility (Fig 5A&B). In all of the lines tested there were many individual females that had no progeny as well as females with varying distributions of progeny. In some of the lines (*w*Mel2a, *w*MelCS2a, *w*MelCS2b) there were outlier females who had large rescue of fertility, some in the range of what we observe for wildtype *bam* fertility (Fig 4B&C). However, for most of the individuals tested, fertility was not fully rescued, e.g., restored to wildtype levels (Fig 4B&C, Fig 5A&B). Although the largest effect of *W. pipientis* on *bam^L255F^*/*bam^null^* rescue was amongst the *w*MelCS-like variants, there were other notable variants outside of the *w*MelCS clade that showed high increase in female fertility, particularly *w*Mel2a. Overall these data further indicate a genetic interaction between *bam* and *W. pipientis* that is specific from the overall effect of *W. pipientis* on female fertility.

**Fig 5.**
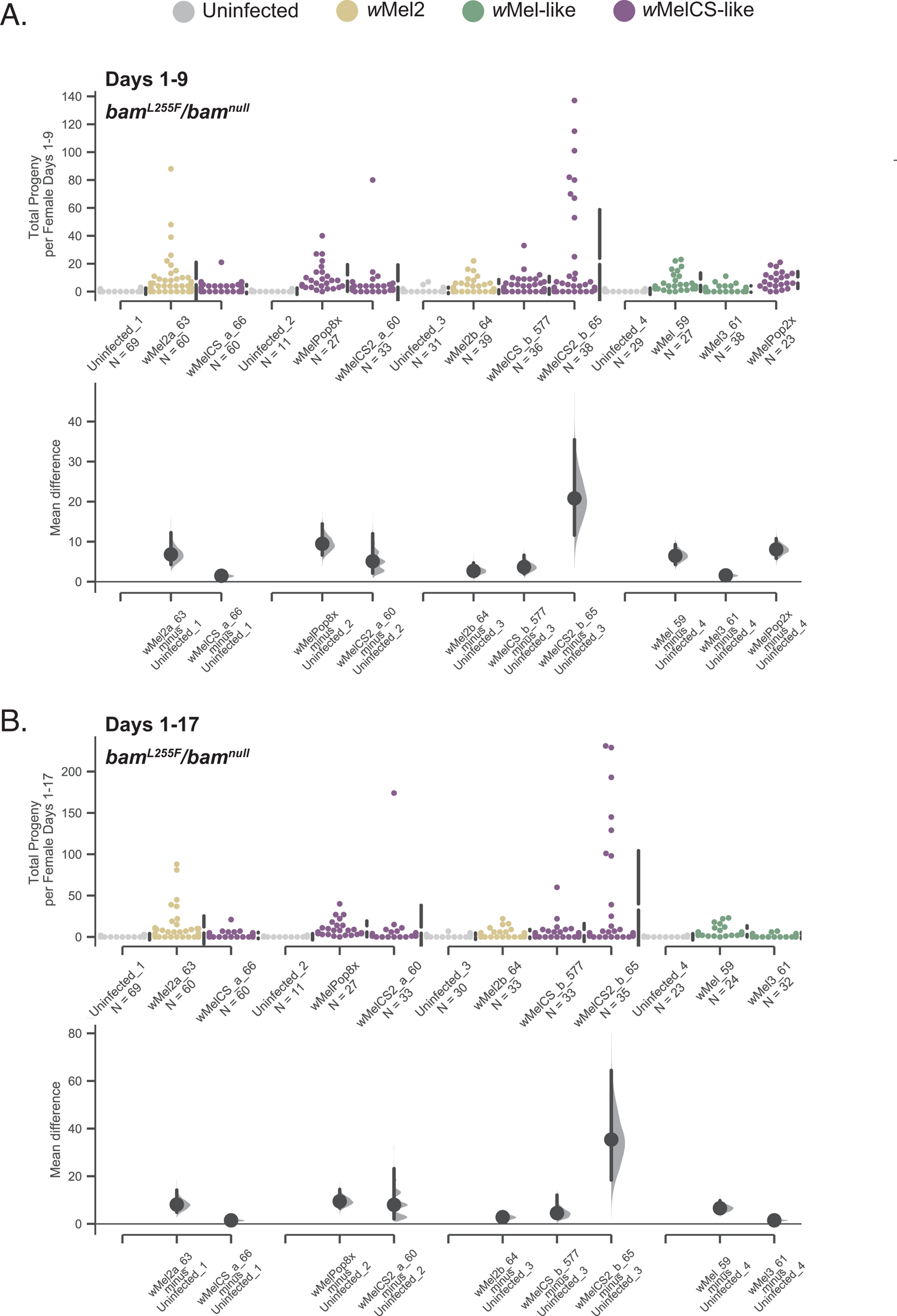
Total progeny per female and mean difference of progeny per female for the *w^1118^*; *bam^L255F^*/*bam^null^* genotype infected with ten *W. pipientis w*Mel variants compared to uninfected. (A) Swarm and Cumming estimation plots showing the total progeny per female for each *W. pipientis* infected line counted on days 1-9 post eclosion. As the fertility assays were performed in batches, each batch is compared to its own uninfected control. All *W. pipientis* variants were associated with significant increases in female fertility compared to uninfected controls (bootstrap 95% confidence interval, effect size). (B) Swarm and Cumming estimation plots showing the total progeny per female for each *W. pipientis* infected line counted on days 1-17 post eclosion. As the fertility assays were performed in batches, each batch is compared to its own uninfected control. All *W. pipientis* variants were associated with significant increases in female fertility compared to uninfected controls (bootstrap 95% confidence interval, effect size). *w*MelPop2X is not reported for this time frame, as most females died after day 9.

### All *w*Mel variants rescue the ovarian *bam* hypomorph cytological defect but to varying degrees

Although we wanted to assess the rescue of the *bam* hypomorphic fertility phenotype by *W. pipientis* because we can also compare this to the effect of *W. pipientis* on wildtype *bam* fertility, we also wanted to know if genetic variation in *W. pipientis* affected the rescue of the *bam* hypomorphic cytological phenotype. Because fertility is affected by environmental and genetic differences that act in later developmental stages outside of *bam*’s function early in gametogenesis, we also measured the effect of *W. pipientis* variation on the rescue of the cytological *bam* mutant phenotype. Since the partial loss of function of *bam* results in over-proliferation of GSCs and reduced differentiation, we can quantify *bam* function by assaying for differentiated germ cells. The cytological *bam* hypomorphic phenotype in females consists of tumorous ovarioles filled with GSC-like cells, with few properly developing egg chambers (Fig 6A). Properly developing egg chambers contain polyploid nurse cells, which feature large, easily identifiable nuclei (Fig 6A, arrowheads). We counted the number of nurse cell positive egg chambers as a measure of GSC daughter differentiation and thus *bam* function. We found that all of the *W. pipientis* variants tested significantly increased the number of nurse cell positive egg chambers, and that *W. pipientis* variants in the *w*MelCS-like clade showed highest numbers of nurse cell positive egg chambers (Fig 7B). We then pooled together the data individually for *w*Mel-like, *w*Mel2-like, and *w*MelCS-like and asked if there was a difference in mean number of cysts containing nurse cells per ovary between each clade (Fig 6C-D). We found that *w*MelCS-like variants had a significantly higher effect size than *w*Mel-like and *w*Mel2-like (Fig6C&D), and that wMel2-like had a higher effect than *w*Mel-like (Fig 6E). These results are consistent with the fertility assay results for the *bam^L255F^*/*bam*^null^ hypomorph genotype, however we observed a more obvious trend of higher *bam* rescue by the *w*MelCS-like variants in this assay. All of the *W. pipientis* variants with the highest effect size of increased *bam* function are in the *w*MelCS-like clade. It is unclear whether this higher rescue is due to titer or another genetic difference between these clades, but these data indicate that genetic variation in *W. pipientis* impacts the *bam* rescue phenotype.

**Fig 6.**
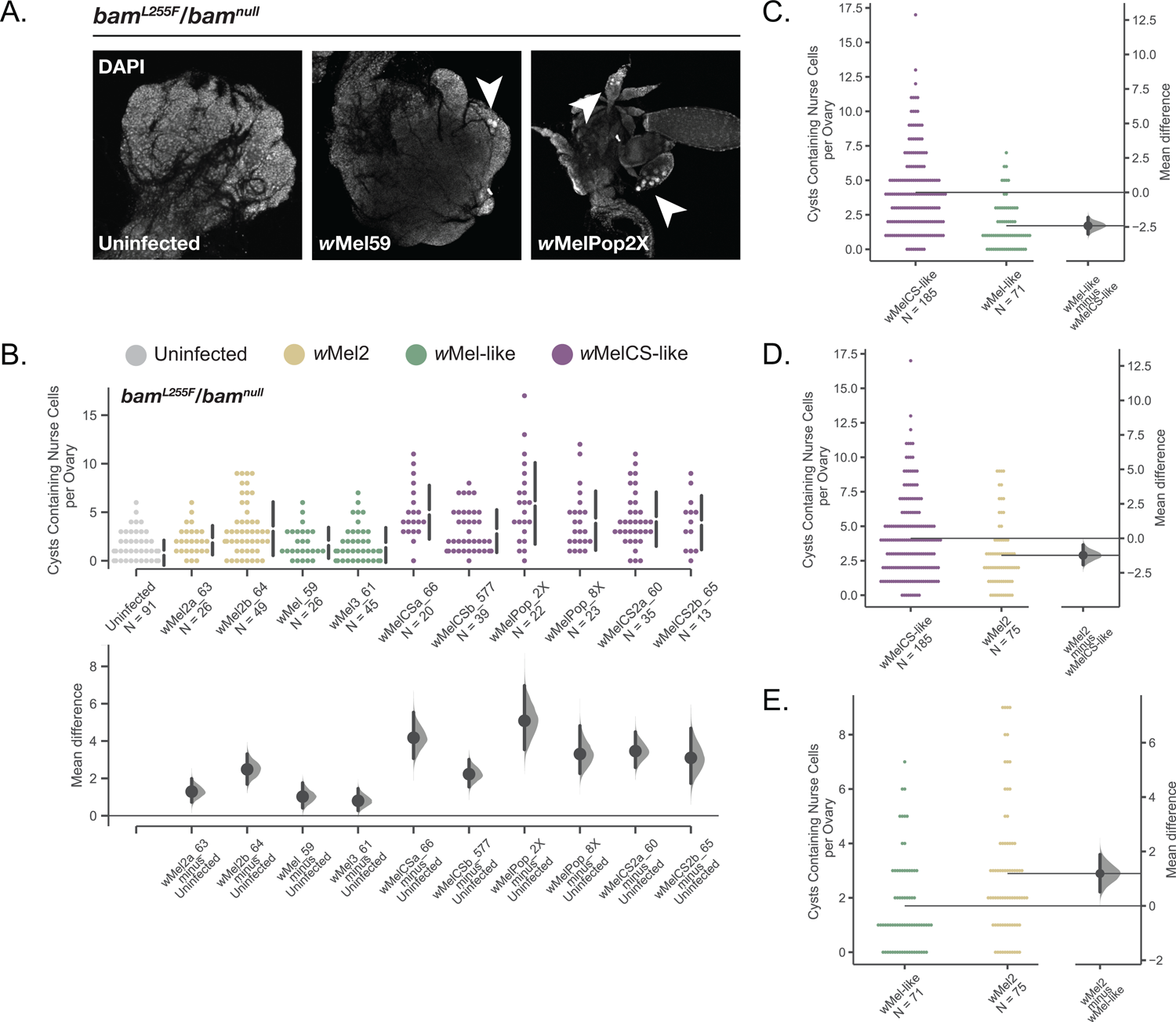
Total cysts containing nurse cells per ovary and mean difference of nurse cell positive cysts for the *w^1118^*; *bam^L255F^*/*bam^null^* genotype infected with ten *W. pipientis* wMel variants compared to uninfected. (A) Representative images of *bam^L255F^*/*bam^null^* ovaries from uninfected, *w*Mel59, and *w*Melpop2X infected females. Nurse cell positive cysts labeled with arrowheads. (B) Swarm and Cumming estimation plots showing the total cysts containing nurse cells per ovary for each *W. pipientis* infected line assayed on 2-3 day old females. All *W. pipientis* variants are associated with a significant increase in nurse cell positive cysts per ovary (bootstrap 95% confidence interval, effect size). *W. pipientis* variants in the *w*MelCS-like clade have the highest effect on nurse cell positive cysts. (C-E) Swarm and Gardner-Altman plots showing pairwise comparisons of nurse cell positive cyst counts pooled together for each *w*Mel clade. *w*MelCS-like variants exhibit higher effects on nurse cell positive cysts compared to *w*Mel2 and *w*Mel-like variants (95% confidence interval, effect size), and *w*Mel2 variants exhibit a higher effect on nurse cell positive cysts than *w*Mel-like (95% confidence interval, effect size).

**Fig 7.**
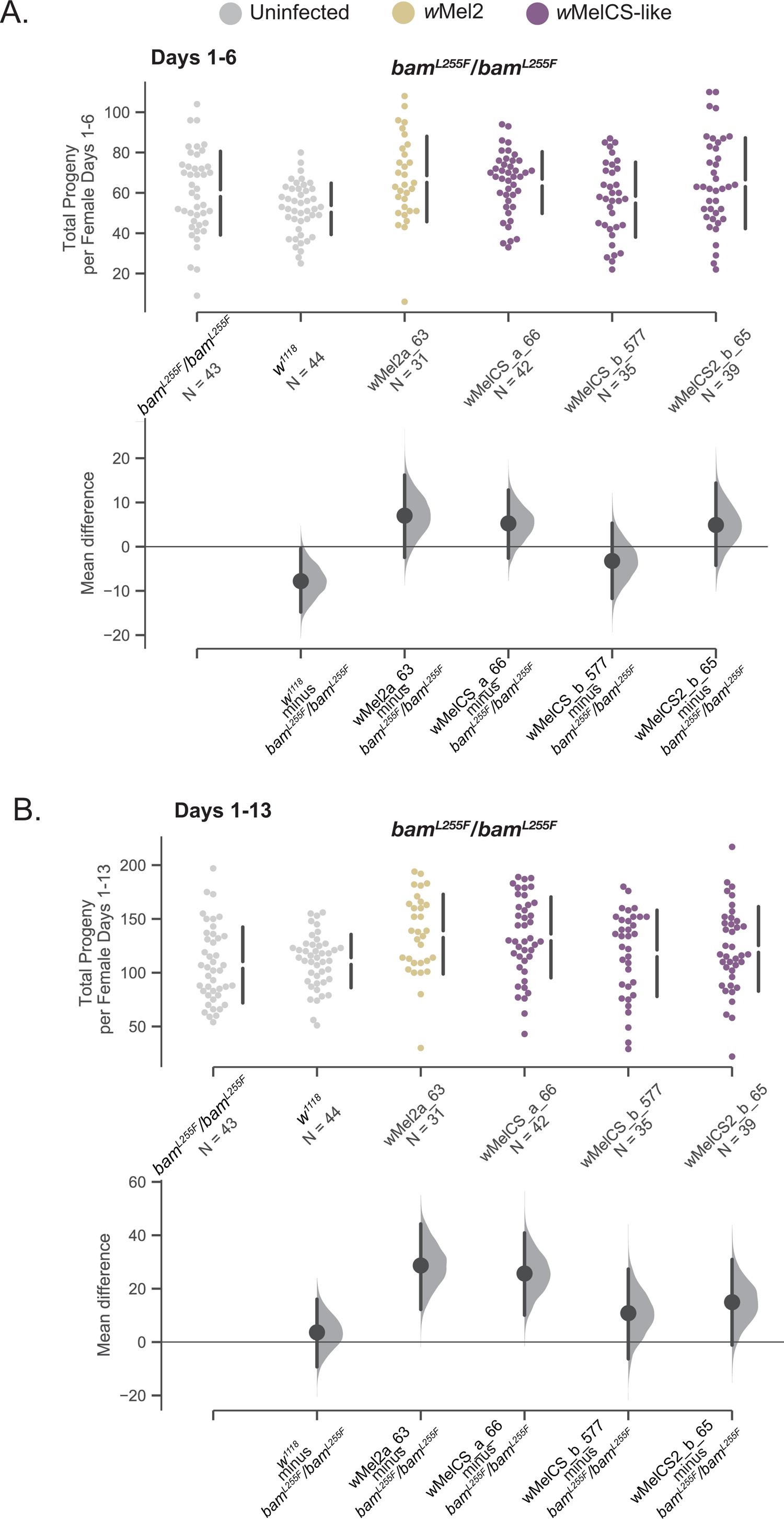
Total progeny per female and mean difference of progeny per female for the *w^1118^*; *bam^L255F^*/*bam^L255F^* genotype infected with four representative *W. pipientis* variants compared to the uninfected control. (A) Swarm and Cumming estimation plots showing the effect of *Wolbachia* variants on *bam^L255F^*/*bam^L255F^* female fertility over days 1-6 of eclosion. We included uninfected *bam*^+^/*bam*^+^ females as well to compare *bam^L255F^*/*bam^L255F^* fertility to *bam*^+^/*bam*^+^ fertility. *W. pipientis* variants did not have differential effects on female *bam^L255F^*/*bam^L255F^* fertility during this time frame. The *bam^L255F^*/*bam^L255F^* genotype regardless of *W. pipientis* infection showed a wider range of progeny compared to *bam*^+^/*bam*^+^ females, and uninfected *bam*^+^/*bam*^+^ mean fertility was significantly lower than uninfected *bam^L255F^*/*bam^L255F^* mean fertility (95% confidence interval, effect size). (B) Swarm and Cumming estimation plots showing the effect of *W. pipientis* variants on *bam^L255F^*/*bam^L255F^* female fertility over days 1-13 of eclosion. *W. pipientis* variants had differential effects on female *bam^L255F^*/*bam^L255F^* fertility. No variants had a negative effect on fertility. *Wolbachia* variants wMel2a_63, wMelCS_a_66 showed significant positive effects on fertility. The *bam^L255F^*/*bam^L255F^* genotype regardless of *W. pipientis* infection showed a wider range of progeny compared to *bam*^+^/*bam*^+^ females, but uninfected *bam*^+^/*bam*^+^ and uninfected *bam^L255F^*/*bam^L255F^* mean fertility was not significantly different.

To ask if the nurse cell positive egg chambers resulted in oocytes capable of development if fertilized, and therefore represent properly differentiated GSC daughters, we picked two representative *W. pipientis* variants of high and low rescue and counted the total number of eggs laid per female over days 1-3. In *w^1118^*; *bam*^+^/*bam*^+^ females, neither *W. pipientis* variant affected the number of eggs laid per female (Fig S1A). In contrast to the *w^1118^*; *bam*^+^/*bam*^+^ females and consistent with the nurse cell assay, we observed few eggs laid by uninfected *w^1118^*; *bam^L255F^*/*bam^null^* females, with significantly more eggs produced by the lower rescuing *W. pipientis w*Mel59, and the highest number of eggs laid by females infected with the high rescuing line, *w*MelCSa_66 (Fig S1B).

### Homozygous *bam^L255F^* females exhibit more variable fertility which is not negatively affected by *W. pipientis*

In light of the variable effect of infection by some *W. pipientis* variants on wildtype *bam* fertility after day 9, and that the uninfected *bam^L255F^*/*bam^L255F^* females did not exhibit a statistically significant fertility defect (Fig 2G, Fig 4B&C), we asked if infection by a subset of the *W. pipientis* variants we assessed in the wildtype *bam* background had any effect on the fertility of *bam^L255F^*/*bam^L255F^* females. We chose three *W. pipientis* variants that did not have a significant effect on wildtype *bam* female fertility for the entire time frame (*w*Mel2a_63, *w*MelCSa_66, and *w*MelCSb_577 Fig 7A&B), and one that had a significant negative effect on female fertility in the second half of the experiment (*w*MelCS2b, Fig 7B). We measured fertility of homozygous females infected with *W. pipientis* variants as previously described. In both *bam*^+^/*bam*^+^ and *bam^L255F^*/*bam^L255F^* backgrounds, no *w*Mel variants affected fertility during the first half of the experiment (Fig 4B, Fig 7A). However, in contrast to the *bam*^+^/*bam*^+^ background where *w*MelCS2b negatively affected female fertility across all days measured, we found that in the *bam^L255F^*/*bam^L255F^* background *w*MelCS2b did not have a significant effect on female fertility (Fig 7B). Additionally, in contrast to our findings in the *bam*^+^/*bam*^+^ background that *w*Mel2a_63 and *w*MelCSa_66 did not significantly affect female fertility, we found that in the *bam^L255F^*/*bam^L255F^* background both of these variants had a significantly positive effect on female fertility across all days measured (Fig 7B).

Overall, we observed a more positive effect of *W. pipientis* on female fertility in the *bam^L255F^*/*bam^L255F^* background than in the *bam*^+^/*bam*^+^ background. As the differential effects of *W. pipientis* on female fertility in both *bam* backgrounds were only in older females, this may be related to the general decline in fecundity as the female ages. Although we did not observe a tumorous ovarian phenotype in the *bam^L255F^*/*bam^L255F^* females, it is possible that *W. pipientis* is interacting with a subtle *bam* phenotype that becomes more apparent as the fly ages and oogenesis declines. This further indicates that there is an effect of *bam* genotype and *W. pipientis* genotype on the interaction between *W. pipientis* and female fertility, and that *W. pipientis* does not just generally increase female fertility in a wildtype *bam* background. Additionally, this interaction does not require the *bam* tumor phenotype.

Interestingly, during days 1-6 uninfected wildtype *bam* females had significantly fewer progeny than uninfected homozygous *bam^L255F^* females (Fig 7A). We also observed that the total progeny per female of the homozygous *bam^L255F^* genotype had a wider range than wildtype, which was not impacted by *W. pipientis* and is therefore likely a subtle *bam* mutant phenotype (Fig 7A). As *bam* regulates GSC daughter differentiation, its mis-regulation may increase or decrease fertility.

## Discussion

### Revisiting the phenotypes of classic *bam* alleles using CRISPR/Cas9 in a controlled genetic background

We previously reported that infection with *w*Mel rescued the fertility defect of *bam^BW^/bam^Δ59^* hypomorphic mutant females (Flores et al. 2015). We sought to better understand this interaction for multiple reasons. The first reason is that we hypothesize that an interaction between *W. pipientis* and *bam* may be driving the adaptive sequence divergence of *bam*. However, in order to understand if and how this interaction could be adaptive, we need to understand the nature of the manipulation of *bam* function by *W. pipientis*. Second, *W. pipientis* is of immense interest as a potential means to control vectors of some human diseases (Moreira et al. 2009).

One such way to find the *W. pipientis* loci necessary for the interaction between bacteria and the host would be to systemically mutagenize the bacteria and then screen for variants that enhance or reduce the phenotype of interest. However, *W. pipientis* are obligate endosymbionts that cannot be cultured, and so are currently not amenable to such genetic manipulation. While some studies have identified *W. pipientis* loci that interact with *D. melanogaster* by transgenically expressing *W. pipientis* loci in *D. melanogaster*, this requires identifying candidate *W. pipientis* loci that affect the host phenotype of interest (Ote et al. 2016; Le Page et al. 2017). An alternative method to identify candidate *W. pipientis* loci that affect the host phenotype is to use genetic variation in *W. pipientis* to test for differences in the effect of *W. pipientis* variants on the host phenotype.

Chrostek et al. (Chrostek et al. 2013) generated isogenic *Drosophila* lines infected with previously described and genetically distinct *w*Mel variants of *W. pipientis*. The original *bam^BW^* mutant was not in this host genetic background and thus we would not be able to conclude that any differential interaction between *bam* and these variants was due to *W. pipientis* variation and not host genetic variation. Therefore, we used CRISPR/Cas9 to create the same amino acid mutant present in the original *bam^BW^* hypomorph allele in the *w^1118^* isogenic background. To ensure we had the tools to assess the *W. pipientis* rescue phenotype as previously described, we also generated a new *bam* null allele using CRISPR/Cas9 in the same *w^1118^* genetic background.

We have now confirmed that these two new alleles behave similarly to the alleles we previously used to document the interaction between *w*Mel and *bam*. Of note, our *bam^L255F^/bam^null^* hypomorph exhibits a stronger fertility defect and GSC daughter differentiation defect in comparison to our previous findings in the *bam^BW^/bam^Δ59^* hypomorph (Flores et al. 2015). Here, for uninfected *bam^L255F^/bam^null^* females, we observed a mean progeny per female of ∼2 and we observed a mean of ∼2 cysts containing nurse cells per ovary (Fig 5, Fig 6). For the *bam^BW^/bam^Δ59^* hypomorph, we previously observed a mean progeny per female of ∼40 and ∼2 nurse cell containing cysts per ovariole (Flores et al. 2015). Notably, each ovary contains 15-20 ovarioles, so 2 nurse cell containing cysts per ovariole would correspond to ∼30-40 nurse cell containing cysts per ovary. Therefore, *w*Mel not only rescues the mild GSC daughter differentiation defect in the original *bam^BW^/bam^Δ59^* hypomorph, but *w*Mel and the other nine variants tested here also rescue the stronger *bam* GSC daughter differentiation defect we observe in the *bam^L255F^/bam^null^* hypomorph. This is especially notable, since *w*Mel does not rescue fertility or GSC daughter differentiation in *bam* null mutants studied by Flores et al. (2015). Therefore, the *w*Mel rescue of the *bam* mutant phenotype requires some functional *bam* gene product, but is also robust to a more severe fertility and differentiation defect. The difference in severity between the two *bam* hypomorphic genotypes also highlights the importance in controlling genetic background to rigorously assess fecundity phenotypes. There have been a handful of studies assessing the phenotypes of diverse *W. pipientis* in isogenic backgrounds (Chrostek et al. 2013; Chrostek et al. 2014; Chrostek and Teixeira 2018; Gruntenko et al. 2019), however there have been no documented studies of interactions between *Drosophila* mutants and *W. pipientis* in these backgrounds. Recent work on the interaction between *W. pipientis* and *Sex lethal* in the female germline showed that *W. pipientis* rescues the loss of GSCs in some *Sxl* mutants through a *nanos* dependent interaction with the *W. pipientis* protein TomO (Ote et al. 2016). However, when researchers transgenically expressed *TomO* in *Sxl* hypomorphic ovaries, the GSC number was rescued, but fertility was not. This result indicated that there are other mechanisms *W. pipientis* is using to fully rescue GSC number and fertility in *Sxl* mutants. We believe the strategy we describe here could be used to further define genetic interactions such as the *Sxl* and *W. pipientis* interaction (Starr and Cline 2002). As we and others have also had success with CRISPR/Cas9 in non-*melanogaster* species, we believe this type of systematic analysis could be extended to assess interactions of host genes and *W. pipientis* in other species outside of *Drosophila melanogaster*.

### *w*Mel *W. pipientis* variants do not broadly increase fertility in the *w*^1118^ genetic background

Due to the shared *w*^1118^ genetic background of the *W. pipientis* infected lines, we were able to rigorously assess the effect of different *W. pipientis* variants on wildtype *bam* female fertility. In addressing if and how *W. pipientis* may drive the adaptive evolution of *bam*, it was important for us to know if *W. pipientis* has an effect on fertility in a wildtype *bam* background. Here, we are not measuring specifically how *W. pipientis* is modulating a functional *bam* allele, as fertility is affected by many loci, but if we observed a large effect of a particular *W. pipientis* variant on fertility, this would motivate further experiments to assess functional *bam* activity in lines infected with that *W. pipientis* variant. We observed no line with increased fertility for any of the *W. pipientis* variants, indicating that *W. pipientis* is not generally increasing fertility through *bam* or another pathway in this genetic background. However, we also observed that some *W. pipientis* variants had a negative effect on female fertility as the females aged. We cannot distinguish whether this effect is due to *W. pipientis* mis-regulating the germline or some other developmental consequence of its infection. In fact, the lines with the highest reported titers in males showed the largest negative impact on female fertility (Chrostek et al. 2013). Additionally, Serga et. al 2014 reported lower fecundity for females from a natural population in Uman, Ukraine infected with *w*MelCS compared to females infected with *w*Mel. These observations highlight the complexity of the interaction between *W. pipientis* and *Drosophila*, where depending on the genetic background of the host, the fitness effect of *W. pipientis* on a phenotype may vary. This effect could further vary based on aspects of the host’s environment where the manipulation of fertility by *W. pipientis* may not always be beneficial. Here we may expect the host and microbe to evolve ways of evading each other, which may become more or less apparent in different genetic backgrounds. Of note, our observations were made under laboratory conditions in a highly inbred line, and so we cannot be sure these *W. pipientis* variants impart the same effects on female fertility in natural populations.

### *w*Mel *W. pipientis* variants across three clades genetically interact with *bam*

We previously showed that a single *w*Mel variants partially rescued the fertility and GSC daughter differentiation defect of the *bam^BW^/bam^Δ59^* hypomorph. Here we asked if there was variation in the *bam* rescue phenotype among *W. pipientis variants*. The ten variants we used have been fully sequenced and differ by varying degrees of sequence divergence. All ten *W. pipientis* variants of the three clades tested (III, VI and VIII; see Fig 4A) rescued the fertility defect and the cytological defect of the *bam^L255F^*/*bam^null^* hypomorph. Therefore, none of the variants tested contained genetic variation that suppressed the interaction between *bam* and *W. pipientis*. We did find that in both the counts of nurse cell positive egg chambers and the fertility assays, *w*MelCS-like *W. pipientis* variants (clade VI) showed the highest rescue (Fig 5, Fig 6). This pattern was the clearest in the counts of nurse cell positive egg chambers as expected, since the presence of nurse cell positive egg chambers reflects GSC daughter differentiation, a phenotype that is a more direct output of *bam* activity. It is likely that the higher level of variability in the fertility assays is also due to the variable effect the *W. pipientis* variants have on other stages of development post GSC daughter differentiation. However, we feel it is important to also measure the effect of *W. pipientis* on fertility, since we cannot assess the count of nurse cell positive egg chambers for *bam* alleles that do not show a tumorous mutant phenotype, and it gives us insight into how *W. pipientis* infection could be adaptive. For example, would there be a fitness tradeoff between a high fecundity (GSC daughter differentiation) rescue and a low fertility (adult progeny) rescue? Measuring the impact of *W. pipientis* on multiple stages of oogenesis and reproduction therefore gives us insight into any further complexities in the interaction between GSC genes and *W. pipientis*.

We found that the *w*MelCS-like variants had the highest rescue effect on *bam* function, another example of a *W. pipientis* induced *D. melanogaster* fitness phenotype of which *w*MelCS-like variants exhibit the highest effects. The *w*MelCS-like variants tested here also confer the highest levels of viral resistance to *D. melanogaster* males (Chrostek et al. 2013). Additionally, *w*MelCS-like variants have a stronger effect on the thermal preference of the *D. melanogaster* host, with *w*MelCS-like variants conferring preference for cooler temperatures compared to *w*Mel-like and uninfected *D. melanogaster* (Truitt et al. 2019). Another study showed that *w*MelCS-like variants increased stress resistance in *D. melanogaster,* and no observed effect from *w*Mel-like variants (Gruntenko et al. 2017).

*w*MelPop is a virulent derivative of *w*MelCSb and characterized by uncontrolled proliferation and titer caused by increased copy number of the Octomom locus. *w*MelPop2X and *w*MelPop8X exhibit 2X copies of the Octomom locus and anywhere from 4X-8X copies, respectively (Chrostek and Teixeira 2018).The high rescue of both the cytological and fertility defect of the *bam^L255F^*/*bam^null^* by *w*MelPop2X is suggestive that titer and possibly other phenotypes that increase interactions with the host are contributing to the rescue of *bam*. We observed lower rescue for *w*MelPop8X, indicating that the increasing copy number of the Octomom locus that causes apoptosis and early death likely negatively affects host fertility (Zhukova and Kiseleva 2012; Chrostek and Teixeira 2018). *w*MelPop2X is closely related to *w*MelCSb, with the only variation between them being a synonymous SNP and the amplification of the octomom locus which contains *W. pipientis* loci predicted to be involved with nucleic acid binding, and thus likely how it increases its titer, as well as proteins with predicted homology to eukaryotic domains (Chrostek and Teixeira 2018). These loci may then increase the interaction of *W. pipientis* and its host (López-Madrigal and Duarte 2020). A natural next step would be to determine the *W. pipientis* factors that manipulate GSC daughter differentiation.

### Possible adaptive interactions between *W. pipientis* and *bam*

It is well established that disrupting *bam* function negatively effects fertility (McKearin and Spradling 1990; Ohlstein and McKearin 1997; Lavoie et al. 1999; Flores et al. 2015), and *bam^L255F^*/*bam^null^* females are almost completely sterile. However, we do not predict such deleterious alleles to reach high frequency in natural populations. In fact, we have not found this nucleotide variant segregating in any of the natural populations of *D. melanogaster* that have been sampled in the *Drosophila* genome Nexus (Lack et al. 2015). Thus, while studying this mutant is effective in further refining how *bam* and *W. pipientis* interact, we cannot conclude that this type of interaction occurs in natural populations. However, given the severity of the mutant and strength of *W. pipientis*’s rescue, one hypothesis is that if *W. pipientis* is increasing GSC daughter differentiation when it is not favorable for the host, the host would evolve a way to evade *W. pipientis*’s manipulation of this pathway. In the case of our lab generated mutant, it is possible that when *bam* is not fully functional that reproduction is more sensitive to manipulation by *W. pipientis*. *W. pipientis* have been documented to respond to changes in the host environment, as *W. pipientis* gene expression is affected by host age and sex, and as *W. pipientis* transmission requires functional oogenesis, it reasonable to hypothesize that *W. pipientis* is sensitive to changes in gametogenesis (Rice et al. 2017; Newton and Sheehan 2018; Russell et al. 2020).

An additional observation we made is that *bam^L255F^*/*bam^L255F^* females show a broader range of adult progeny per female compared to wildtype *bam* females regardless of *W. pipientis* infection status. The individual *bam^L255F^*/*bam^L255F^* females exhibit both higher and lower than average fertility, indicating that mis-regulating *bam’s* differentiation function could both increase and decrease fertility. This phenotype is worth further investigation, since although *bam* shows a signature of positive selection, we do not know specifically what aspect of *bam* function is adaptive. If variation in *bam* function can both increase and decrease mean fertility, and *W. pipientis* has a generally positive effect on fertility of *bam^L255F^*/*bam^L255F^* females, this is further evidence that *W. pipientis* may be able to manipulate *bam* in order to increase oogenesis for its own benefit, and that genetic variation at *bam* could affect the regulation of oogenesis. Therefore, if *W. pipientis* increased the rate of oogenesis to ensure its own transmission and this was not favorable for the host, *bam* may be in conflict with *W. pipientis* to regulate oogenesis in a favorable manner for the host. Here we see that although the average female fertility of uninfected *bam^L255F^*/*bam^L255F^* mutants is not significantly different from wildtype, there is a wider distribution of individual female fertility, indicating that the *bam* hypomorphic phenotype is an increased variance in fertility (Fig 7). Since these females are in a common genetic background and we do not expect genetic variation between individual females, the *bam*-mediated mis-regulation of differentiation may set off a cascade of other genetic mis-regulation that results in higher or lower fertility over time. We see that infection by some *W. pipientis* variants in this background increases female fertility, indicating that perhaps *W. pipientis* can restore the mis-regulation of GSC daughter differentiation, and even increase reproductive output through this mechanism. It would be interesting to use this genotype for future experiments to explore the possibility that this *bam* mutation may disrupt the mechanism *D. melanogaster* evolved to evade *W. pipientis*’s effect on differentiation.

An interesting question that remains is how *W. pipientis* have evolved in their interaction with GSC genes, including *bam*. While *w*MelCS-like and *w*Mel-like variants share a most recent common ancestor about 8000 years ago, *w*MelCS-like variants were recently (∼2000 years ago) replaced by *w*Mel-like variants in natural populations of *D. melanogaster* (Riegler et al. 2005; Richardson et al. 2012). However, this replacement has not been complete, and there are global populations still infected with *w*MelCS variants (Riegler et al. 2005; Nunes et al. 2008). Interestingly we find that the *w*MelCS variants we assayed show higher rescue of *bam^L255F^*/*bam^null^* female fertility and nurse cells. If *bam* and *W. pipientis* are evolving in an arms race, it could be that the evolutionarily more recent *w*Mel-like variants have not evolved the same level of interaction with *bam* as *w*MelCS has. Future work to investigate these dynamics should include sampling populations that are still infected with *w*MelCS as well as those infected with *w*Mel and asking if there is any evidence of genetic differentiation at *bam*. Some populations have been identified that are still infected with *w*MelCS, such as a natural population of *D. melanogaster* from Uman, Ukraine that has been infected with both *w*Mel and *w*MelCS and has been monitored yearly for infection frequency, some additional Paleartic populations, and a population in the Netherlands (Early and Clark 2013; Bykov et al. 2019; Serga et al. 2021).

Additionally, we could utilize existing *W. pipientis* sequence variation from natural populations to ask if there is any evidence of associations between *bam* variation and *W. pipientis* variation. If we were to sample populations that are differentially infected with *w*Mel and *w*MelCS to assess genetic differentiation at *bam* as discussed above, we could also do the same for *W. pipientis* variants. One caveat being that if we did not observe genetic signatures of adaptive evolution in *W. pipientis* loci, this does not mean that *W. pipientis* and *bam* are not coevolving, as we do not know the true infection history of a *Drosophila* population with *W. pipientis* and thus which *W. pipientis* variant may have been in conflict with *bam.* Additionally, there may be general *W. pipientis* functions that are interacting with *bam* and not a single locus (e.g., loci that regulate titer).

Further work should focus on determining the *W. pipientis* loci contributing to the different magnitude of *bam* rescue. However, as has been previously pointed out by Chrostek et al. (Chrostek et al. 2013), between the *w*Mel clades and *w*MelCS clades there are eight indels and 108 SNPs, including differences in the coding sequence of 58 genes. So further work would have to be done to narrow down which variants contribute to the degree of rescue. A next step could be moving out to more divergent *W. pipientis wMel* variants (for example, *w*Au that infects *D. simulans* (Miller and Riegler 2006)) to ask how recently the interaction with *bam* evolved. To complement this, it would be beneficial to perform these same rescue experiments with *bam* hypomorphs in *D. simulans*.

## Acknowledgements

This work was supported by National Institutes of Health grant R01-GM095793 to C.F.A. We would like to thank Luis Teixeira for his advice on assessing the effect of *W. pipientis* variation on *Drosophila* fertility and for the gifts of the *w^1118^* isogenic lines infected with *W. pipientis* variants used in this study. We also would like to thank Kristen Rose Baxter for her contributions to our early fertility experiments. We are also grateful to Mariana F. Wolfner, Andrew G Clark, Miwa Wenzel and Catherine Kagemann for valuable input and discussion of these experiments and results and helpful comments on this manuscript.

---

**Fig S1.**
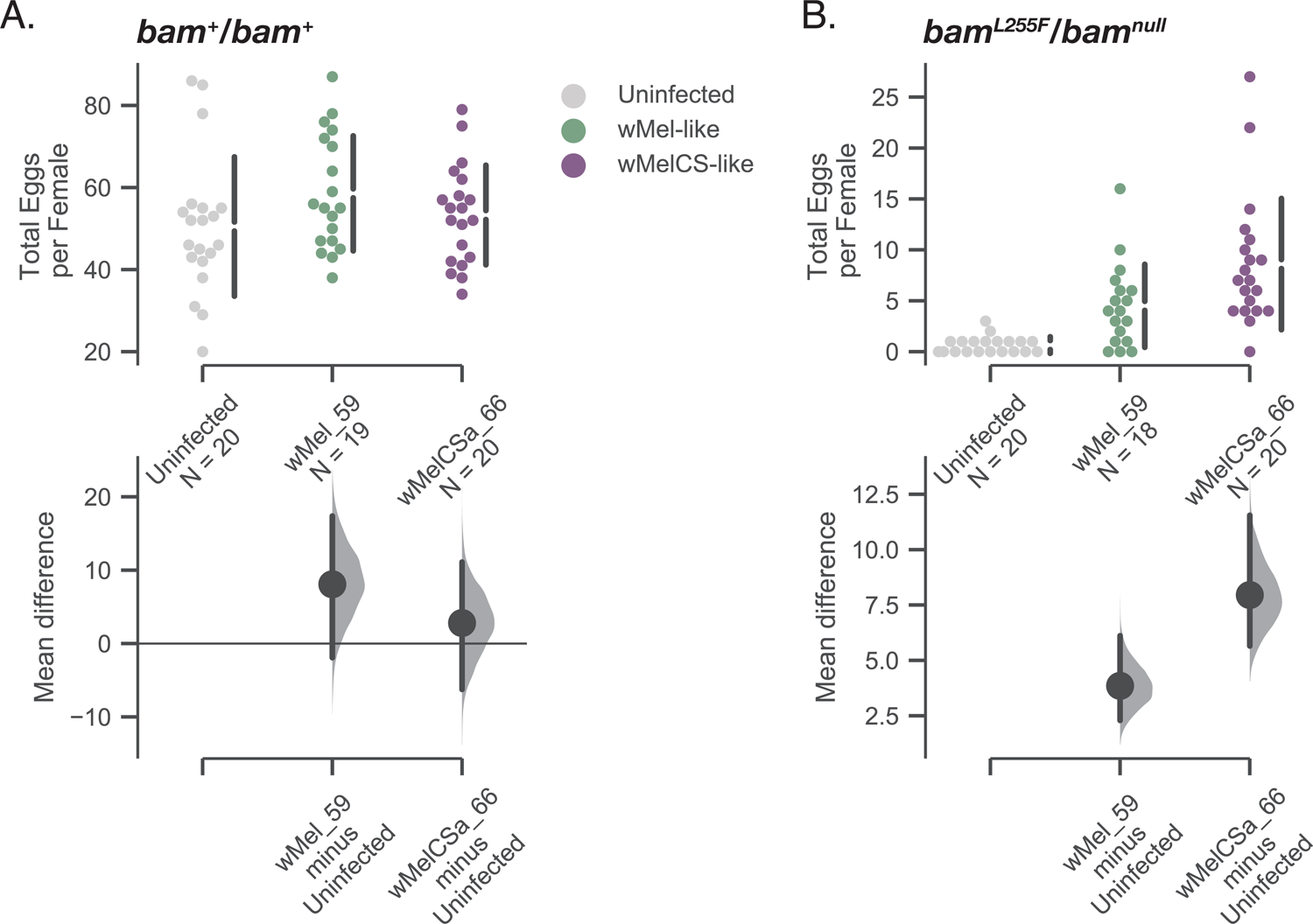
Total eggs per female counted over 3 days for the *w^1118^*; *bam*^+^/*bam*^+^ genotype and the *w^1118^*; *bam^L255F^*/*bam^null^* genotype infected with representative *W. pipientis* variants from the *w*Mel-like and *w*MelCS-like clades. (A) Swarm and Cumming estimation plots showing the total eggs per female for *w^1118^*; *bam*^+^/*bam*^+^ lines infected with *w*Mel59 or *w*MelCSa66. Infection with either variant does not significantly affect total eggs per female. (B) Swarm and Cumming estimation plots showing the total eggs per female for *w^1118^*; *bam^L255F^*/*bam^null^* females infected with the same two *W. pipientis* variants. Infection with either *w*Mel_59 or *w*MelCSa_66 significantly rescues the number of eggs laid per female compared to the uninfected control (95% confidence interval, effect size). Consistent with the results of the nurse cell assay, *w*MelCSa_66 shows a higher rescue effect than *w*Mel_59.

